# Experimental testing of two urban stressors on freshwater biofilms

**DOI:** 10.1101/2023.06.11.544504

**Authors:** Romain VRBA, Isabelle LAVOIE, Nicolas CREUSOT, Mélissa EON, Débora MILLAN-NAVARRO, Agnès FEURTET-MAZEL, Nicolas MAZZELLA, Aurélie MOREIRA, Dolors PLANAS, Soizic MORIN

## Abstract

Aquatic ecosystems and their communities are exposed to numerous stressors of various natures (chemical and physical), which impacts are often poorly documented. In urban areas, the use of biocides such as dodecyldimethylbenzylammonium chloride (DDBAC) and their subsequent release in wastewater result in their transfer to urban aquatic ecosystems. DDBAC is known to be toxic to most aquatic organisms. Artificial light at night (ALAN) is another stressor that is increasing globally, especially in urban areas. ALAN may have a negative impact on photosynthetic cycles of periphytic biofilms, which in turn may result in changes in their metabolic functioning. Moreover, studies suggest that exposure to artificial light could increase the biocidal effect of DDBAC on biofilms. The present study investigates the individual and combined effects of DDBAC and/or ALAN on the functioning and structure of photosynthetic biofilms. We exposed biofilms in artificial channels to a nominal concentration of 30 mg.L^-1^ of DDBAC and/or ALAN for 10 days. ALAN modified DDBAC exposure, decreasing concentrations in the water but not accumulation in biofilms. DDBAC had negative impacts on biofilm functioning and structure. Photosynthetic activity was inhibited by > 90% after 2 days of exposure, compared to the controls, and did not recover over the duration of the experiment. Biofilm composition was also impacted, with a marked decrease in green algae and the disappearance of microfauna under DDBAC exposure. The integrity of algal cells was compromised where DDBAC exposure altered the chloroplasts and chlorophyll content. Impacts on autotrophs were also observed through a shift in lipid profiles, in particular a strong decrease in glycolipid content was noted. We found no significant interactive effect of ALAN and DDBAC on the studied endpoints.

**Graphical abstract:** 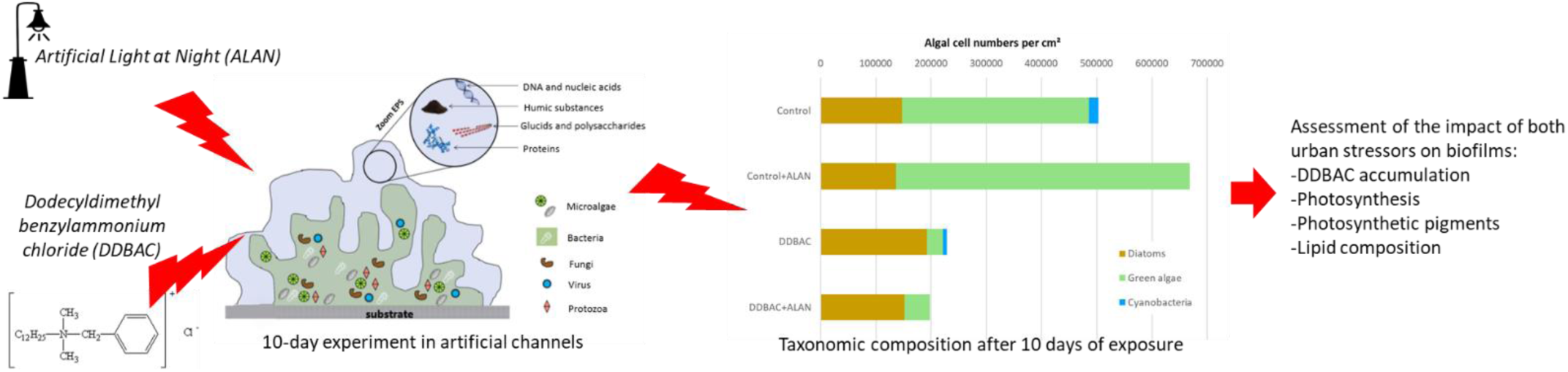

## 1- Introduction

Biocides are widely used in different areas of activity including agriculture and industry, as well as in a multitude of domestic products (e.g. pharmaceutical, personal care and household products) (Abbott et al., 2020). Among these biocides, dodecyldimethylbenzylammonium chloride (DDBAC), a quaternary ammonium compound, is commonly used as an active substance in medical disinfection products, as well as in household products such as detergents. Recently, following the regulation of some broadly used disinfectant agents such as triclosan, quaternary ammonium compounds like DDBAC have become popular substitutes (Sreevidya et al., 2018). Moreover, DDBAC and all other benzalkonium chloride derivatives have been shown to be very effective against the SARS-CoV virus (Rabenau et al., 2005), resulting in their widespread use as disinfectants during the Covid 19 pandemic (US EPA, 2020). This multi-purpose product eventually reaches wastewaters, where its concentration varies greatly depending on the land use, population and wastewater treatment facilities, among other factors. For example, a concentration of 6 mg.L^-1^ was recorded in hospital effluents (Kümmerer et al., 1997) due to the intensive use of DDBAC as a disinfectant. In municipal wastewater effluents, DDBAC concentrations were observed to range from just a few ng per liter up to 170 µg.L^-1^ (Clara et al., 2007; Martínez-Carballo et al., 2007; Zhang et al., 2015).

Once in the aquatic environment, DDBAC is relatively stable to photodegradation and its half-life in surface waters can reach 180 days, although the presence of photosensitizers can reduce this half-life to a week (US EPA, 2006). The organic carbon-water partition coefficient (Koc) of DDBAC is 5.43 and, therefore, it has a high adsorption ratio on sewage sludge, sediments or humic substances (van Wijk et al., 2009). Although the transfer of DDBAC to urban aquatic ecosystems has been recognized, the ecotoxicity of DDBAC is poorly documented. The toxicity of DDBAC to cells is due to its quaternary ammonium polar head, which has the capacity to bind to the surface membrane, while the alkyl lipophilic chain alters the phospholipid bilayer. This alteration can rapidly lead to membrane disruption and progressive lysis of the cell (Eich et al., 2000). Regarding aquatic organisms, there is redundancy in the biological models used for toxicity assessment where most studies have focused on *Daphnia magna* to determine the acute toxicity of DDBAC (Kreuzinger et al., 2007; Leal et al., 1994; Chen et al., 2014; Lavorgna et al., 2016). To date, *D. magna* is the most sensitive organism to DDBAC, where the concentration needed to immobilize 50% of organisms (EC50) was of 5.9 µg.L^-1^ (US EPA, 2006). As a result, the US EPA established a non-observed adverse effect concentration (NOAEC) of 4.15 µg.L^-1^ for aquatic invertebrates. Phytoplanktonic organisms seem to be less sensitive to DDBAC than aquatic invertebrates. Indeed, other studies investigated the effects of DDBAC on various microalgae species and the range of EC50 based on growth was found to vary between 58 µg.L^-1^ for *Skeletonema costatum* (Kreuzinger et al., 2007) to 203 µg.L^-1^ for *Chlorella vulgaris* (Sütterlin et al., 2008). However, to our knowledge, the chronic effects of DDBAC exposure on natural aquatic communities has not be considered.

In this study, we focused on stream biofilms (periphyton), which are complex structures housing autotrophic microorganisms (e.g. cyanobacteria, green algae, diatoms), bacteria, fungi, and other heterotrophic organisms such as microfauna (Battin et al., 2016). Biofilms are ubiquitous in aquatic environments, including urban streams and ponds. They form the basis of aquatic food webs, providing a source of energy for primary producers, including essential fatty acids (Brett and Müller-Navarra, 1997). Biofilms can also be found downstream of wastewater treatment plants (WWTPs) and have been used as ecological indicators of stress (Tamminen et al., 2022; Tlili et al., 2020). The early detection of ecosystem impairment using autotrophic biofilms can be based on functional (e.g., photosynthesis) and structural endpoints describing biodiversity based on taxonomy as well as pigments or lipid profiles (Sabater et al., 2007; Morin & Artigas, in press). Biofilms are also efficient bioaccumulators of organic substances carried by waters under dissolved or particulate forms (Bonnineau et al., 2021). Given the toxic mode of action of DDBAC, impacts on biofilm fatty acids, particularly on membrane phospholipids, are likely to occur. The attack on phospholipid membrane may also result in the disruption of key functions of biofilm such as photosynthesis. For example, Pozo-Antonio and Sanmartin (2018) showed a significant decrease in chlorophyll *a* and photosynthetic efficiency of phototrophic biofilms from church walls after a DDBAC treatment. The effect of DDBAC as a biofouling removal agent was significantly enhanced when combined with artificial light or UV irradiation.

Artificial light at night (ALAN) has become a global pollution concern and, as of 2014, more than 20% of the world’s land surfaces between 75°N and 60°S were exposed to a light-polluted sky (Falchi et al., 2016). Most urban areas have developed along rivers and coastlines, increasing the exposure of urban aquatic environments to ALAN. Concerns about the impact of ALAN on aquatic ecosystems and research on this topic are quite recent (Perkin et al., 2011). Indeed, ALAN was only recognized as harmful for freshwater ecosystems in the early 2000s (Longcore and Rich, 2004). ALAN could lead to disruptions in nycthemeral cycles of autotrophic organisms in biofilms by inducing variability in the maximum efficiency of photosynthesis (Maggi and Serôdio, 2020). ALAN can also alter taxonomic composition by favouring certain autotrophic groups over others, thereby exerting differential selection, as shown in a study where cyanobacteria proportions in biofilm decreased under ALAN (Grubisic et al., 2017).

The objective of this study was to determine the individual and combined effects of two anthropogenic stressors typical from urban aquatic ecosystems (DDBAC and ALAN) on autotrophic organisms within biofilms, using structural and functional descriptors of impact. To this aim, we performed a 10-day experiment in which natural autotrophic biofilms were exposed in laboratory channels to DDBAC and/or ALAN. We hypothesized that, given the Koc of the substance, DDBAC would bioaccumulate in the biofilms, generating toxicity to microorganisms. Functional (photosynthetic efficiency) and structural (taxonomic composition, pigment and lipid profiles) descriptors were predicted to be impacted by DDBAC exposure. ALAN was expected to modify the development of microalgae, thus to impact the proportions of algal groups, to alter photosynthetic efficiency to a certain extent, and to increase the sensitivity of microalgae to additional stress. As a consequence, we hypothesized that co-exposure to ALAN and DDBAC would enhance the toxic effects of the biocide.

## 2- Material and methods

### 2.1- Experimental design

Biofilms were grown in a DDBAC-free pond in Cestas, near Bordeaux, France. A previous study classified this small water body as a hypereutrophic pond (Chaumet et al., 2019; Neury-Ormanni et al., 2020). Glass slides (∼200 cm²) were immersed in the photic zone, at a depth of 30-50 cm. Colonization took place in winter (December 2020 to April 2021), during five months to collect sufficient amounts of biomass. At this period, daylight irradiance on the top surface on the water (5 cm deep) was below 100 µmol.s^-1^.m^-^². At the end of the colonization period, 12 glass slides covered by mature biofilms were randomly distributed among four series of experimental units (EU) for the experiment. The EUs consisted of experimental channels of about 30 L filled with pond water that had been previously filtered (20 µm) to remove suspended material and most planktonic organisms. The slides were divided among three parallel channels per EU, connected to 10-L tanks, for subsequent exposure of the biofilms to the different treatments (DDBAC and ALAN). Biofilms corresponding to day 0 (d0, 5 slides) were collected immediately after their recovery from the pond.

On d0, two EUs were contaminated with a solution of dodecyldimethylbenzylammonium chloride (DDBAC; Sigma Aldrich, France; CAS: 139-07-1, purity: >99%) to reach a concentration of 30 mg.L^-1^. This concentration was selected to elicit an effect on photochemical efficiency while using as low a concentration as possible, based on the observed EC5 of preliminary dose-response experiments (Appendix A). In order to assess only the effect of altered photoperiod, not light level, a non-stressful incident light intensity of 20 µmol.s^-1^.m^-^² was selected. Light was kept on overnight and then from day 1 (d1), one control series and one DDBAC series were exposed to an alternating 14 h day / 10 h night photoperiod (alternated photoperiod, AP) and the other two series (one control, one DDBAC) were exposed to a 24 h day photoperiod (continuous photoperiod, CP) throughout the course of the experiment (Figure 1). Room temperature was maintained constant at 20.5 ± 0.1°C, while water temperature was kept at 18.7 ± 0.2°C. The tanks were refilled with 3 L of filtered pond water contaminated at 30 mg.L^-1^ DDBAC on day 4 (d4) to compensate for water evaporation.

**Figure 1:**
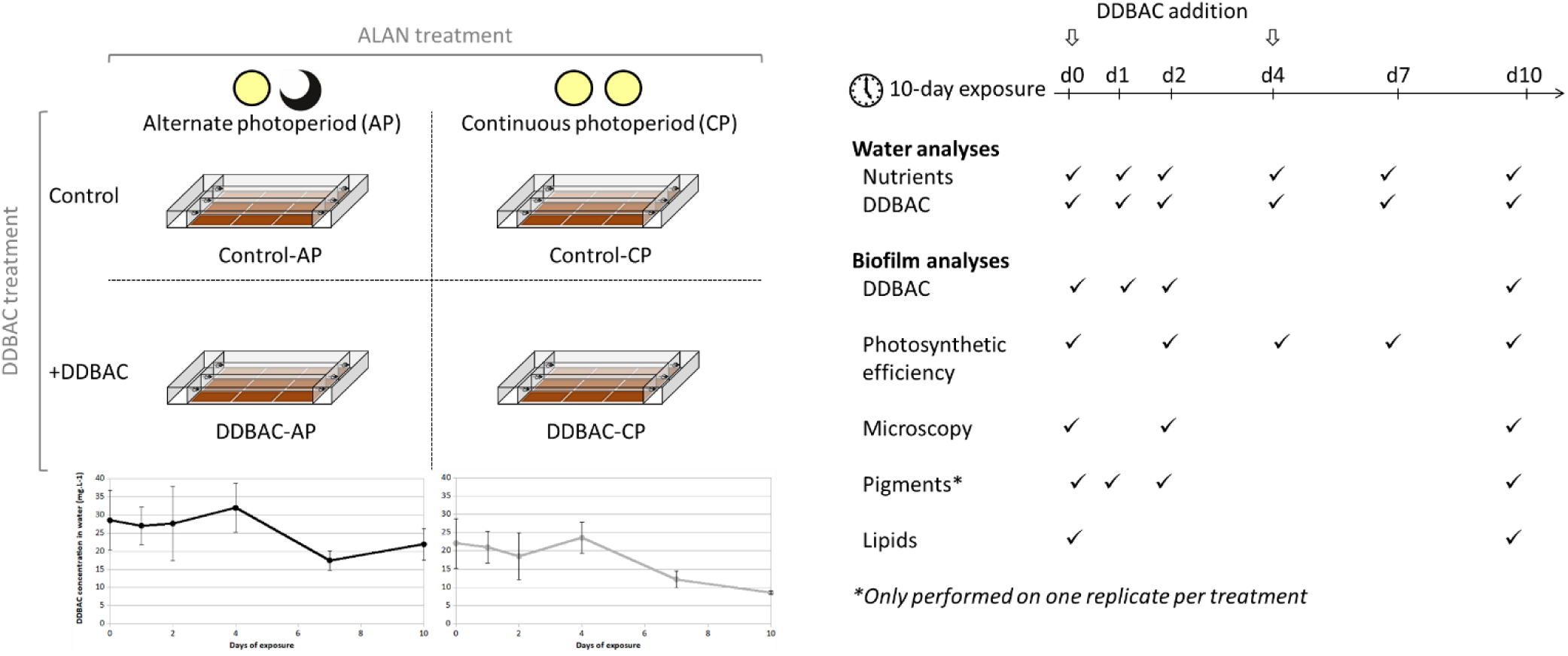
Experimental design and sampling strategy.

### 2.2- Water chemistry measurements

On d0, d1, d2, d4, d7 and d10, 20 mL water samples were taken from all channels, filtered on 1-μm PTFE filters and stored at 4°C in the dark until analysis (performed within 48 h of collection). Nutrients and mineral salts were analysed as described in Chaumet et al. (2019), using a Metrohm 881 Compact Ionic Chromatograph pro (Metrohm). Anion analysis (PO4^-^, NO3^-^, NO2^-^, Cl^-^ and SO4^2-^) was performed using a Supp 4/5 Guard/4.0 precolumn followed by a Metrosep A Supp5 – 250/4.0 column. The mobile phase was a mixture of a solution of 3.2 mmol.L^-1^ Na2CO3 and a solution of 1 mmol.L^-1^ NaHCO3. Cation analysis (Na^+^, K^+^, Ca^2+^, Mg^2+^ and NH^4+^) was performed using a C4 Guard/4.0 precolumn followed by a Metrosep C6 -250/4.0 column). All precolumns and columns were provided by Metrohm. The eluent used was a mixture of 2.5 mmol.L^-1^ HNO3 and a solution of 1.7 mmol.L^-1^ 10,12-Pentacosadynoic acid (PCDA). The limits of quantification (LOQ) of the different ions analysed are shown in Table 1.

**Table 1.** Principal ions and DDBAC concentrations (mg.L^-1^) in the four experimental treatments along the ten days of exposure. Results are expressed as the mean and standard error for the whole experiment. Two-way ANOVAs were conducted, with ALAN and DDBAC as the two factors, except for DDBAC concentrations (One-way ANOVA assessing the effect of ALAN). DDBAC = contaminated biofilm; Control = non-exposed biofilm; AP = alternated photoperiod; CP = continuous photoperiod; LOQ = Limit of quantification.

| Treatment | DDBAC | NO<sub>2</sub><sup>-</sup> | NO<sub>3</sub><sup>-</sup> | PO<sub>4</sub><sup>3-</sup> | SO<sub>4</sub><sup>2-</sup> | Cl<sup>-</sup> | NH<sub>4</sub><sup>+</sup> | Na<sup>+</sup> | K<sup>+</sup> | Ca<sup>2+</sup> | Mg<sup>2+</sup> |
| --- | --- | --- | --- | --- | --- | --- | --- | --- | --- | --- | --- |
| Control-AP | <LOQ | 0.04 ± 0.03 | 1.69 ± 0.27 | 0.03 ± 0.02 | 21.0 ± 1.8 | 36.4 ± 2.7 | 0.09 ± 0.05 | 22.0 ± 1.9 | 4.80 ± 0.43 | 57.8 ± 5.1 | 4.90 ± 0.42 |
| DDBAC-AP | 26.6 ± 2.5 | 0.02 ± 0.00 | 0.66 ± 0.17 | 0.03 ± 0.02 | 18.9 ± 0.8 | 35.5 ± 1.4 | 0.11 ± 0.05 | 19.7 ± 0.9 | 4.34 ± 0.20 | 52.2 ± 2.5 | 4.41 ± 0.20 |
| Control-CP | <LOQ | 0.02 ± 0.01 | 1.35 ± 0.23 | 0.01 ± 0.00 | 17.5 ± 0.3 | 30.3 ± 0.4 | 0.09 ± 0.06 | 18.5 ± 0.3 | 4.12 ± 0.06 | 48.4 ± 0.6 | 4.09 ± 0.06 |
| DDBAC-CP | 18.5 ± 2.0 | 0.02 ± 0.00 | 0.75 ± 0.10 | 0.03 ± 0.02 | 17.8 ± 0.3 | 33.0 ± 0.6 | 0.10 ± 0.06 | 18.8 ± 0.9 | 4.21 ± 0.11 | 49.5 ± 1.2 | 4.18 ± 0.09 |
| LOQ | 0.001 | 0.005 | 0.01 | 0.01 | 0.005 | 0.01 | 0.005 | 0.01 | 0.0025 | 0.15 | 0.15 |
| Significant factor(s) and interaction(s) | ALAN | None | DDBAC | NA | DDBAC | DDBAC | NA | DDBAC | NA | DDBAC | DDBAC |
| p | <0.001 | >0.05 | <0.001 | NA | <0.05 | <0.01 | NA | <0.05 | NA | <0.05 | <0.05 |

DDBAC concentrations in the water were monitored frequently over the experiment, at d0, d1, d2, d4, d7 and d10. Three samples of 20 mL were collected from each channel and stored at −20°C together with the stock solution until analysis. The samples were analysed using an Ultimate 3000 HPLC coupled with an API 2000 triple quadrupole mass spectrometer. We used a Gemini® NX-C18 column from Phenomenex as a stationary phase. The mobile phase was 90:10 5 mM ammonium acetate/acetonitrile. We worked in isocratic mode, so the composition of the mobile phase was constant during the analysis. The flow rate was set at 0.6 mL.min^-1^ and the injection volume was set at 20 µL. An internal standard of benzyl-2,3,4,5,6-d5-dimethyl-n-dodecylammonium chloride (DDBAC-d5) was used (Cluzeau, France; CAS: 139-07-1, purity: >98%) at an equivalent of 100 µg.L^-1^ in both samples and standard solutions for calibration. The mass spectrometer was operated in multiple reaction monitoring (MRM) mode with a total cycle time of 0.8 seconds. The MRM transitions were 304>91 (quantitation) and 304>212 (confirmation) for DDBAC. For DDBAC-d5, the MRM transitions were 309>96 and 309>212. The declustering potential was set to 65 V and the collision energies were set either to 45 or 30 V for the quantitation or confirmation transition, respectively. Samples were diluted and the calibration range was from 1 to 200 µg.L^-1^. Quality controls were regularly injected at concentrations of 5 and 25 µg.L^-1^, as well as analytical blanks.

### 2.3- Biofilm biological endpoints

#### 2.3.1- Photosynthetic efficiency

We sampled 2 cm² of biofilm from three different glass slides for each treatment on d0, d2, d4, d7, and d10, corresponding to three pseudoreplicates per treatment per sampling. Each biofilm sample was then suspended in 3 mL of channel water and gently shaken for homogenization. Photosynthetic activity (effective photosystem II quantum yield) was assessed on suspensions of biofilm (Genty et al. 1989) within one hour after collection using a Pulse Amplifitude Modulation fluorimeter (Phyto-PAM, Heinz Walz GmbH, Germany) in quartz cuvettes under continuous agitation (Emitter–Detector Unit PHYTO-ED). Biofilm suspensions were adapted for 15 min to light (164 μmol.m^−2^.s^−1^) before fluorescence measurements at the same active radiation, in Actinic Light mode. The quantum yield values reported are mean values of five measurements per replicate, averaged for the three pigment groups using the device’s reference spectra. After photochemical efficiency measurements, samples were preserved for further microscopic analyses by adding a few drops of Lugol solution then stored in the dark at 4°C.

#### 2.3.2- Microscopic analyses

For microscopic observations, a Nageotte counting slide (Marienfeld, Germany) was used with 125 µL biofilm suspension samples collected on d0, d2 and d10 to determine initial composition as well as to assess early structural changes and chronic effects of ALAN and DDBAC. Observations were made at x400 magnification under an optical microscope (Olympus BX51) equipped with a digital camera as described in Morin et al. (2010). Ten fields of view were scanned for enumeration of the main taxonomic groups observable on preserved material: diatoms, green algae, cyanobacteria and microfauna. Solitary and colonial algae and cyanobacteria were recorded, and counted as cell numbers to allow for density comparisons. Microfauna organisms were recorded as individuals. All densities were expressed as a function of colonized biofilm surface (i.e., per cm²). Live (intact cell content) and dead (empty frustules) diatoms were counted separately to estimate mortality (Morin et al. 2010).

#### 2.3.3- Analyses from freeze-dried biofilms

Biofilms were scraped from glass slides on d0, d1, d2 and d10 (4 slides per treatment) and were frozen in liquid nitrogen to prevent lipid degradation. The samples were then lyophilized for 24 hours, and used to determine lipid content, DDBAC bioaccumulation, and concentrations of photosynthetic pigments. Analyses focused on neutral lipids (storage lipids: triacylglycerides) and on polar lipids. Polar lipids correspond to phospholipids (lipids typical from cytoplasmic membranes: phosphatidylcholine-PC, phosphatidylethanolamine-PE, and phosphatidylglycerol-PG) and to glycolipids (characteristic from chloroplastic membranes: digalactosyldiacylglycerol-DGDG, monogalactosyldiacylglycerol-MGDG, sulfoquinovosyldiacylglycerol-SQDG). Lipids were extracted from the biofilms (20 mg dry weight) using a mixture of MTBE-methanol (3:1 %v/v) and UPW-methanol (3:1 %v/v) with 150 mg of microbeads. Samples were then homogenized using a FastPrep (MP Biomedicals). The upper organic fraction of the samples (i.e. MTBE) was recovered by centrifugation for lipid analysis, while the lower hydrophilic fraction (mixture of UPW and methanol) was used to determine DDBAC bioaccumulation.

##### 2.3.3.1- Lipid quantification

For lipid analysis, samples from d0 and d10 were evaporated and injection solvent was added before analysis by HPLC-MS/MS following the protocol detailed in Mazzella et al. (in press). Briefly, different stationary and mobile phases were used for the analysis of phospholipids/glycolipids and triglycerides. For the phospholipid and glycolipid analyses, a LUNA® NH2 HILIC column (100 x 2 mm, 3 µm) from Phenomenex was used as the stationary phase, and a mixture of acetonitrile and 40 mM ammonium acetate buffer as the mobile phase. The flow rate was set at 400 µL.min^-1^. The proportions of these two solutions are given in Appendix C. For the analysis of triglycerides and betaine lipids (i.e. diacylglyceryltrimethylhomo-Ser, DGTS), a KINETEX® C8 column (100 x 2.1 mm, 2.6 µm) from Phenomenex was used as the stationary phase, and the mobile phase was a mix of a solution of acetonitrile/water/40 mM ammonium acetate buffer (600/390/10, v/v/v) and a solution of isopropanol/acetonitrile/1 M ammonium acetate buffer (900/90/10, v/v/v). The flow rate was set at 300 µL.min^-1^. The proportions of these two solutions are given in Appendix C. Results were then pre-treated with ANALYST® 1.6.2 software from Sciex. For polar lipids (i.e. glycolipids and phospholipids), the limits of quantification ranged from 0.1 to 0.5 nmol.mg^-1^, depending on the lipid classes. For triglycerides and DGTS, the limits of quantification reached were 0.01 and 0.1 nmol.mg^-1^, respectively. Results were expressed as nmol.g^-1^ freeze-dried biofilm.

##### 2.3.3.2- DDBAC accumulation in biofilms

The same analytical method used for water samples was used to determine bioaccumulated DDBAC in the previously extracted hydrophilic fraction. It was generally necessary to perform significant dilutions (i.e. 10,000-fold) in order to stay within the calibration range. Results were then expressed in the log10 value of the bioconcentration factor (i.e. log(BCF)). The BCF was calculated according to the following formula:

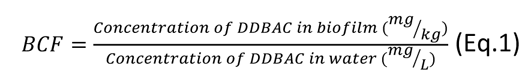

##### 2.3.3.3- Photosynthetic pigments

On selected freeze-dried samples (one replicate per treatment), pigment analyses were also performed following the French standard NF T90-117 (1999). Ten mg of dry biofilm were put in solution using 10 mL of acetone. After 20 minutes of ultrasonication, the mix was filtered on a Büchner filter to remove the solid phase. Absorption was measured at the wavelengths 630 nm, 647 nm, 664 nm, 665 nm and 750 nm with a UV-1800 (Shimadzu) spectrophotometer, and was then remeasured at the same wavelengths after acidification of the samples. The concentrations of chlorophyll pigments and phaeopigments were determined following the equations of Lorenzen (1967) and expressed as µg pigments.mg^-1^ freeze-dried biofilm.

### 2.4- Data analyses

After verification that homogeneity of variances was met, ANOVAs were performed. One-way ANOVA was used to test the effect of ALAN on DDBAC degradation, and two-way ANOVAs were used to assess the individual effect of ALAN and DDBAC, as well as their interaction, on nutrient concentrations. Two-way ANOVAs with sampling dates as repeated measures were applied to assess the individual and interactive effects of ALAN and DDBAC on biofilm endpoints. Verification for homogeneity of variances and normality was performed prior to conducting ANOVAs. Data were processed using R software (R Core Team, 2022).

## 3- Results

### 3.1- Water chemistry

In the control channels, DDBAC concentrations were always below detection limit (Table 1), confirming that no cross-contamination occurred. Contaminant concentrations averaged 26.6 ± 2.5 mg.L^-1^ (AP) and 18.5 ± 2.0 mg.L^-1^ (CP) in the DDBAC treatments. Differences in DDBAC concentrations were observed between the DDBAC treatment channels exposed to an alternated photoperiod and those exposed to a continuous photoperiod (F[1,10] = 232.72; p < 0.001, also see Figure 1), suggesting that degradation occurred under continuous photoperiod. Nutrient concentrations were also modified by DDBAC exposure (Table 1). Indeed, DDBAC addition resulted in lower concentrations of nitrate, sulfate, Cl^-^, Na^+^, Ca^2+^ and Mg^+^ than in the control (Table 1), while ALAN showed no effect on nutrient concentrations.

### 3.2- Bioaccumulation of DDBAC in biofilms

Table 2 shows the log(BCF) for the hydrophilic fraction of biofilm samples at d1, d2 and d10 for AP and CP treatments. DDBAC was recovered from biofilms for every treatment and every sampling time, showing that this compound bioaccumulates well in autotrophic biofilms, with a mean log(BCF) of 3.22 ± 0.25. The amount of DDBAC bioaccumulated in the biofilms showed no significant difference in bioaccumulation between sampling times (F[2,28]= 2.38; p = 0.11) nor ALAN treatments (F[1,28] = 0.28; p = 0.60). Overall, the amount of DDBAC bioaccumulated in biofilms was stable through the experiment and not impacted by ALAN (Time x ALAN interaction: F[2,28] = 0.025; p = 0.98).

**Table 2.** Evolution of DDBAC bioaccumulated in biofilms (expressed in log(BCF)) between the different ALAN treatments (n=3). AP = alternated photoperiod CP = continuous photoperiod.

|  | d1 | d2 | d10 |
| --- | --- | --- | --- |
| AP | $2.18 \pm 1.70$ | $3.03 \pm 0.34$ | $3.40 \pm 0.10$ |
| CP | $3.23 \pm 0.30$ | $2.90 \pm 0.98$ | $3.33 \pm 0.10$ |

### 3.3- Photosynthetic efficiency

Figure 2 shows that photosynthetic activity was stable over the time of the experiment, with a mean value of 0.42 ± 0.01 in the control AP and CP channels, regardless of the ALAN treatment (F[1,42] = 2.65; p > 0.05). Biofilms exposed to DDBAC showed an almost complete inhibition of photosynthetic yield starting from d2 and lasting until the end of the experiment (F[1,42] = 1279; p < 0.001). In the same way as for control channels, photosynthetic activity showed no difference between AP and CP for contaminated channels (F[1,42] = 0.18; p = 0.67).

### Photoperiod; CP = Continuous Photoperiod

### 3.4- Photosynthetic pigments

Chlorophyll *a* concentrations decreased over time in the DDBAC contaminated samples, compared to the control samples (Figure 3). At d2, no chlorophyll *a* was detected in the DDBAC-AP samples, and only phaeopigments were measured. At the end of the experiment, pigment concentrations in DDBAC samples were very low compared to the controls. Unfortunately, the lack of replication does not allow for any conclusion on the effect of the tested treatments on chlorophyll *a* or phaeopigments.

**Figure 2.**
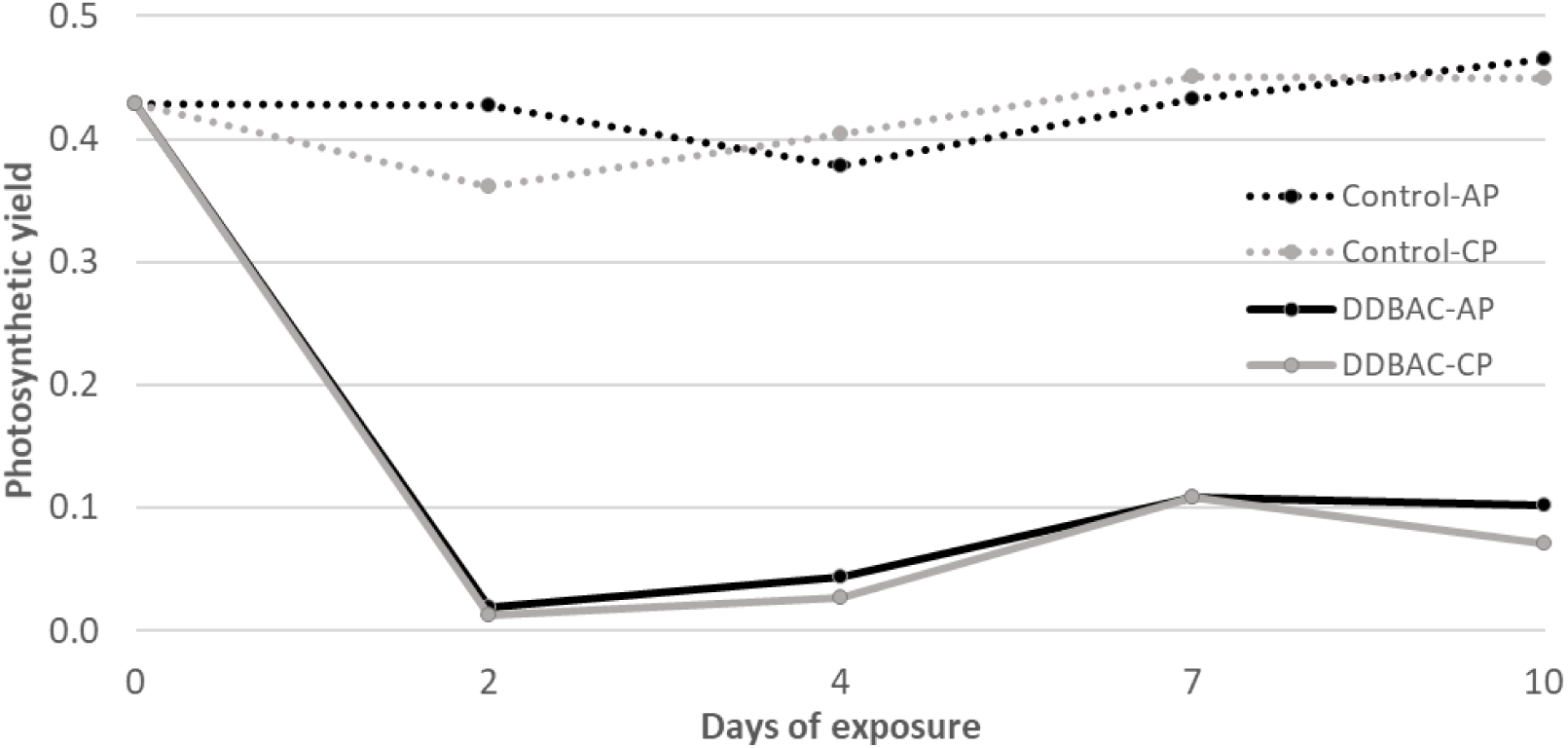
**Evolution of photosynthesis yield in biofilms along the ten days of the experiment (d0: n=5, d2 and d10: n=3). CTRL = non-exposed biofilm; DDBAC = contaminated biofilm; AP = Alternated**

**Figure 3.**
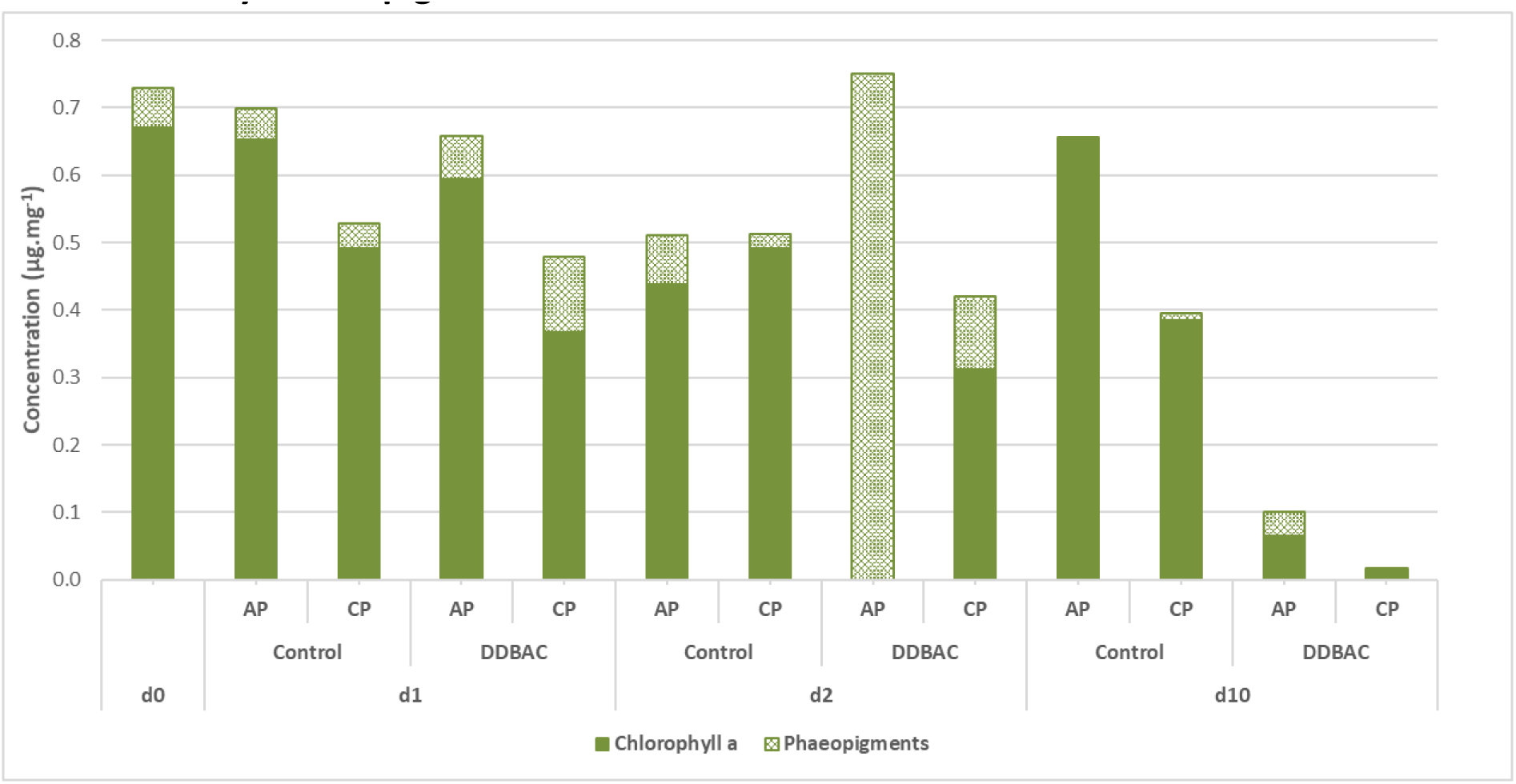
**Changes in photosynthetic pigments content in biofilms along the ten days of the experiment (n = 1). Control = non-exposed biofilm; DDBAC = contaminated biofilm; AP = alternated photoperiod; CP = continuous photoperiod.**

### 3.5- Community composition

Diatom mortality is expressed in %. Results are expressed as mean and standard error for each date and treatment (d0: n=5, d2 and d10: n=3). DDBAC = contaminated biofilm; CTRL = non-exposed biofilm; AP = Alternated Photoperiod; CP = Continuous Photoperiod Diatoms (91%) dominated the biofilm community at d0, whereas low proportions of green algae (5%), cyanobacteria (4%) and microfauna (1%) were observed (Table 3). In the control channels, green algae developed markedly and became the main algal group in terms of density by d10, with no significant effect of the ALAN treatment (e.g. 67% in AP channels and 79% in CP). The density of green algae slightly increased from d0 to d10 in the DDBAC contaminated channels, but not to the same extent as in the control channels. Indeed, a significant Time × DDBAC interaction was highlighted for green algae densities (p < 0.001; Table 3). Microfauna increased in control channels, but had completely disappeared from biofilms exposed to DDBAC by d10. For this component, a significant Time × DDBAC interaction was also noted (p < 0.01; Table 3). ALAN treatments did not have a significant impact on the biofilm taxonomic composition.

**Table 3.** Evolution of the taxonomic composition of biofilms during the experiment. Densities are expressed as individuals×10^3^ / cm², i.e. number of cells for microalgae, and number of organisms for microfauna.

| Date | Treatment | Diatom density | Diatom mortality | Green algae density | Cyanobacteria density | Microfauna density |
| --- | --- | --- | --- | --- | --- | --- |
| d0 | Control | 81.3 ± 33.7 | 7 ± 1 | 4.1 ± 2.3 | 3.8 ± 2.0 | 0.3 ± 0.0 |
| d2 | Control-AP | 141.3 ± 9.1 | 3 ± 0 | 26.7 ± 4.4 | 1.6 ± 2.0 | 0.6 ± 0.3 |
| d2 | DDBAC-AP | 113.2 ± 11.5 | 3 ± 1 | 11.8 ± 2.8 | 0 | 0.2 ± 0.2 |
| d2 | Control-CP | 132.1 ± 68.8 | 3 ± 1 | 51.7 ± 46.0 | 0 | 0.3 ± 0.3 |
| d2 | DDBAC-CP | 145.0 ± 26.1 | 2 ± 1 | 6.2 ± 1.7 | 0 | 0 |
| d10 | Control-AP | 98.2 ± 11.7 | 5 ± 2 | 225.6 ± 133.5 | 11.2 ± 7.8 | 0.9 ± 0.5 |
| d10 | DDBAC-AP | 128.5 ± 26.0 | 2 ± 0 | 18.9 ± 15.0 | 4.8 ± 3.0 | 0 |
| d10 | Control-CP | 91.3 ± 14.2 | 6 ± 1 | 354.2 ± 83.1 | 0 | 1.0 ± 0.2 |
| d10 | DDBAC-CP | 101.0 ± 27.0 | 1 ± 0 | 30.7 ± 8.2 | 0 | 0 |
| Significant factor(s) and interaction(s) | | None | DDBAC | DDBAC × Time | None | DDBAC × Time |
| p |  | >0.05 | <0.05 | <0.001 | >0.05 | <0.01 |

Diatom mortality values (based on the ratio between frustules without cell content and total diatom frustules) significantly decreased under DDBAC exposure (Table 3). In addition, microscopic observations highlighted differences in the aspect (shape, colour) of chloroplasts in diatoms and chlorophytes exposed to DDBAC, becoming highly granular and darker (Figure B.2). These were, however, not considered as ‘dead’ cells in estimating diatom mortality.

### 3.6- Lipid content

On d0, total lipid concentration reached 42.6 nmol.mg^-1^ of biofilm dry weight. Almost half of the lipids were phospholipids, and the other half were glycolipids, while neutral lipids (TAG) represented a minor portion (Table 4). By d10, total lipid concentration had decreased in all treatments down to an average of 23.6 ± 7.53 nmol.mg^-1^ of biofilm dry weight (Table 4). No significant differences were found between AP and CP, while DDBAC significantly affected glycolipids over time (significant Time × DDBAC interaction; p < 0.01). As an example, Figure 4 shows lipid contents under DDBAC and ALAN exposure on d0 and d10. At the beginning of the experiment, phosphatidylglycerol (PG) was the predominant lipid in the biofilms, while phospholipids and glycolipids in the biofilms were evenly distributed. Total lipid amounts decreased over time in the controls, but the proportions of phospholipids and glycolipids remained similar to those at d0, all the main classes of lipids being present. Total phospholipids remained stable over time, whatever the treatment (p > 0.05), but phoshatidylethanolamine (PE) and PG were higher in biofilms exposed to DDBAC compared to controls (Figure 4, Table 4). In the samples exposed to DDBAC, glycolipids such as monogalactosyldiacylglycerol (MGDG) and digalactosyldiacylglycerol (DGDG) decreased significantly, or even became barely detectable, with significant Time × DDBAC interactions (respectively, p < 0.01 and 0.001). Diacylglyceryltrimethylhomo-Ser (DGTS) was never detected and, therefore, is not mentioned further.

**Figure 4.**
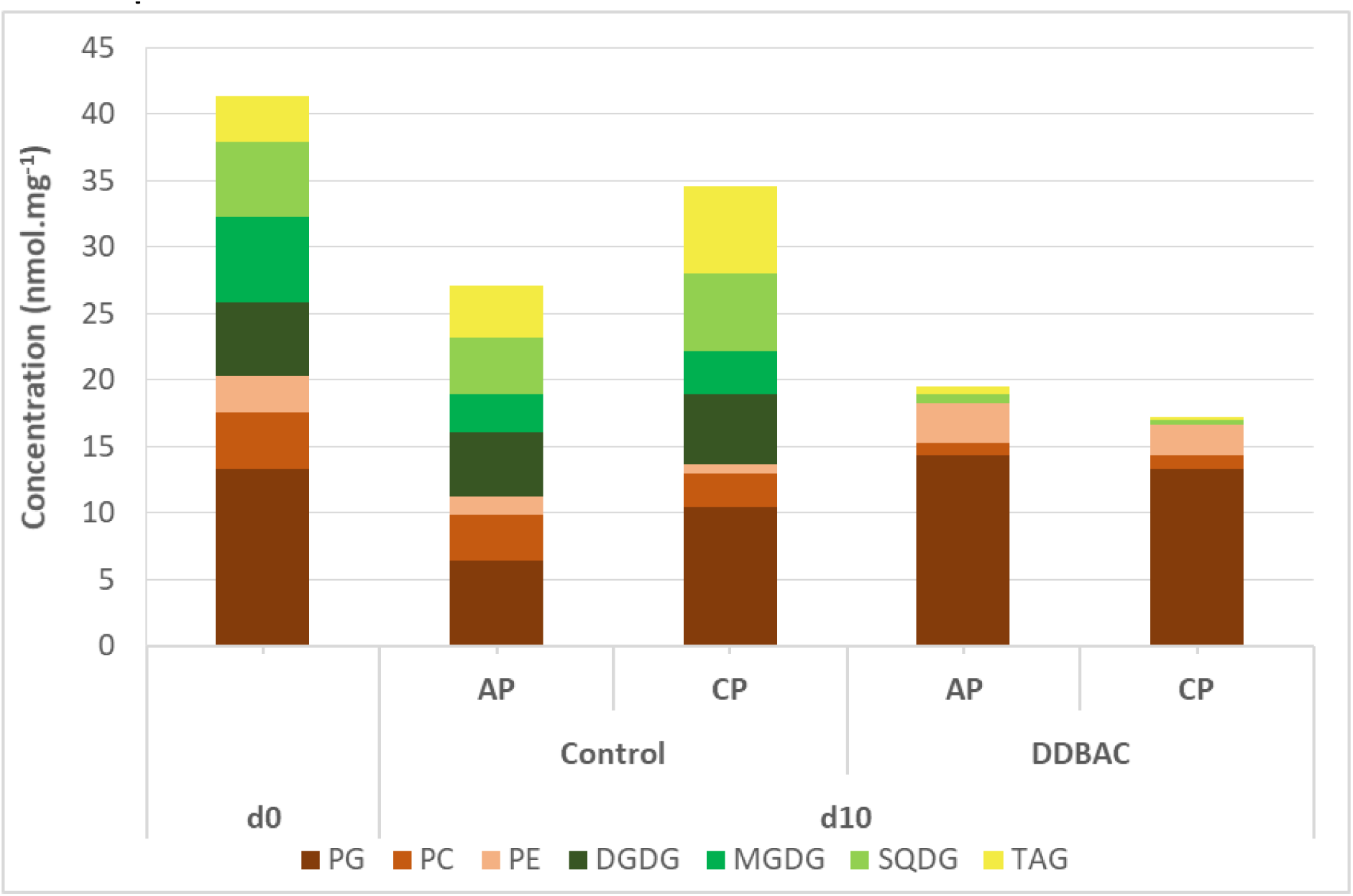
**Effects of DDBAC and continuous photoperiod on the proportion of total lipids and on lipid classes (n=4). DDBAC = contaminated biofilm; CTRL = non-exposed biofilm; AP = alternated photoperiod; CP = continuous photoperiod; PE = phosphatidylethanolamine; PC = phosphatidylcholine; PG = phosphatidylglycerol; DGDG = digalactosyldiacylglycerol; MGDG = monogalactosyldiacylglycerol; SQDG = sulfoquinovosyldiacylglycerol; TAG = triacylglycerides.**

**Table 4:** Evolution of lipid content of biofilms (in nmol.g-1 freeze-dried biofilm) between the beginning and the end of the experiment. Results are expressed as mean and standard error for each date and treatment (n=4). DDBAC = contaminated biofilm; CTRL = non-exposed biofilm; AP = Alternated Photoperiod; CP = Continuous Photoperiod; PE = phosphatidylethanolamine; PC = phosphatidylcholine; PG = phosphatidylglycerol; DGDG = digalactosyldiacylglycerol; MGDG = monogalactosyldiacylglycerol; SQDG = sulfoquinovosyldiacylglycerol; TAG = triacylglycerides. Two-ways ANOVAs on repeated measures were applied on data.

| Date | Treatment | Neutral lipids (TAG) | PG | PE | PC | Σ Phospholipids | DGDG | MGDG | SQDG | Σ Glycolipids | Total lipids |
| --- | --- | --- | --- | --- | --- | --- | --- | --- | --- | --- | --- |
| d0 | Control | 4.2 ± 2.5 | 13.7 ± 3.1 | 2.8 ± 0.5 | 4.6 ± 0.2 | 21.0 ± 3.3 | 5.6 ± 1.2 | 6.4 ± 1.6 | 5.3 ± 1.1 | 17.3 ± 2.4 | 42.6 ± 3.2 |
| d10 | Control-AP | 3.9 ± 2.0 | 6.4 ± 1.0 | 1.3 ± 0.5 | 3.5 ± 1.5 | 11.2 ± 2.4 | 4.8 ± 0.8 | 2.9 ± 0.5 | 4.2 ± 0.7 | 12.0 ± 0.4 | 27.1 ± 4.1 |
| d10 | DDBAC-AP | 0.2 ± 0.3 | 14.3 ± 2.6 | 3.0 ± 0.4 | 0.9 ± 0.4 | 18.2 ± 2.6 | 0 | 0 | 0.7 ± 0.2 | 0.7 ± 0.2 | 19.1 ± 2.6 |
| d10 | Control-CP | 6.6 ± 2.9 | 6.9 ± 6.7 | 0.7 ± 0.1 | 2.6 ± 1.1 | 10.1 ± 7.8 | 5.3 ± 0.2 | 3.3 ± 1.1 | 5.8 ± 1.2 | 14.4 ± 2.2 | 31.1 ± 8.1 |
| d10 | DDBAC-CP | 0.1 ± 0.1 | 13.3 ± 3.5 | 2.3 ± 1.0 | 1.0 ± 1.0 | 16.6 ± 5.2 | 0 | 0 | 0.4 ± 0.2 | 0.4 ± 0.2 | 17.1 ± 5.3 |
|  | **Significant factor(s)** | DDBAC × Time | DDBAC × Time | DDBAC × Time | DDBAC × Time | None | DDBAC 12× Time | DDBAC × Time | DDBAC × Time | DDBAC × Time | DDBAC × Time |
|  | **p** | 0.01 | <0.05 | <0.01 | <0.01 | >0.05 | <0.001 | <0.01 | <0.001 | <0.001 | <0.001 |

## 4. Discussion

### 4.1- Experimental conditions and DDBAC exposure

In this experiment, natural biofilms were transferred to controlled laboratory conditions under ALAN and DDBAC treatments. Changes in environmental conditions were likely to affect biofilm characteristics; control conditions were however favourable to the development of autotrophic biofilms, as highlighted by continuous growth from d0 to d10 and by stable physiological state (assessed through photosynthetic efficiency). During the 10 days, most physicochemical parameters remained stable, and no nutrient depletion was observed at the end of the experiment in any treatment. However, DDBAC seemed to have an indirect effect on the concentrations of certain nutrients over the 10-day exposure period of our experiment. For example, under DDBAC exposure, concentrations of NO ^-^ were lower than in the control (Table 1). DDBAC may have affected the growth of biofilm autotrophs, leaving space for heterotrophic microorganisms consuming nutrients such as nitrates. DDBAC concentrations in the channels under CP decreased over time (Figure 1), and was on average lower than under AP. This could be the result of photodegradation of the contaminant or of an increase in DDBAC retention by biofilms under ALAN conditions. A previous study showed that freshwater biofilms can improve the photodegradation of organic contaminants by being a source of reactive oxygen species (ROS) such as hydroxyl (OH·) or superoxide (O2·) radicals (Yin et al. 2022).

Contrary to the hypothesis of Pozo-Antonio and Sanmartin (2018), who suggested that artificial light could lead to an increase in internalization of the contaminant, we found no significant difference in DDBAC concentrations in biofilm between ALAN treatments (Table 2). Thus, our results do not support the hypothesis of a higher uptake of the substance from the water through increased bioaccumulation under ALAN exposure. This lower concentration might be linked to some bacterial strains such as *Aeromonas hydrophila* that can biodegrade DDBAC and use the degradation product as a source of nitrogen (Patrauchan and Oriel, 2003), converging with the above-mentioned hypothesis related to the decrease in certain nutrients concentrations in the waters under DDBAC treatment. However, resistant strains may not withstand DDBAC concentrations higher than 10 mg.L^-1^ (Kreuzinger et al., 2007).

The potential for DDBAC exposure on biofilm consumers (related to bioaccumulation in the biofilm) was similar between ALAN treatments, independently of differences in DDBAC concentrations in the water. DDBAC accumulation rapidly stabilized to average log(BCF) of 3.22 (Table 2), falling in the range measured for organochlorine substances in biofilms, and greater than previous observations for other pesticides or pharmaceuticals (Bonnineau et al. 2021). Our results demonstrate the retention capacity of biofilms for this compound, suggesting a risk of transfer to primary consumers.

### 4.2- Effects of DDBAC on the biofilm

The first effect of DDBAC observed was the marked and rapid decrease in photosynthetic efficiency (more than 90% compared with the initial yield; Figure 2). This sudden inhibition of photosynthesis was not expected because the exposure concentration was based on the EC5 for photosynthesis inhibition after four hours of exposure determined in a preliminary experiment (EC5 = 30 mg.L^-1^; Figure A). Based on our microscopic observations, diatom mortality was low in all samples (<10%). It should be noted that, following Morin et al. (2010), only diatoms that no longer had chlorophyll in their cell content were considered as dead cells, while cells with chloroplasts were counted as live diatoms. However, many of the cells where chloroplastic content was observed nevertheless seemed to have suffered critical alteration of their photosynthetic equipment (Figure B.2). Their integrity was potentially severely affected, which could explain the loss of photosynthetic yield in biofilms exposed to DDBAC. This deterioration of cell integrity may be due to the fact that DDBAC is able to quickly penetrate the cell (Severina et al. 2001). Despite the lack of replication for pigment analyses, the degradation of chlorophyll pigments in DDBAC treatments (Figure 3) provides additional evidence of the impairment of the algal component of the biofilms that was already highlighted through photosynthetic inhibition and chloroplast alteration.

Marked differences in the autotrophic community composition were highlighted by microscopy counts between d2 and d10 under DDBAC exposure. Indeed, densities of green algae were 10 times lower than under control conditions (Table 3). Despite the lack of replication for pigment analyses, the degradation of chlorophyll pigments in DDBAC treatments (Figure 3) is in line with the impairment of the algal component of the biofilms as highlighted through photosynthetic inhibition and chloroplast alteration. The striking decrease in lipid content in biofilms exposed to DDBAC (Table 4), particularly glycolipids found mainly in thylakoid membranes within photosynthetic cells (Zulu et al., 2018), supports the idea that the contaminant is internalized within phototrophic cells, with deleterious impacts on the autotrophic component. Diatom-rich biofilms are generally source of essential polyunsaturated fatty acids such as eicosapentaenoic acid (Zulu et al., 2018), ensuring a high nutritional quality of this food resource for primary consumers (Brett and Müller-Navarra, 1997). Recently, Demailly et al. (2019) also demonstrated that diatom fatty acid composition can be altered by organic substances such as pesticides. DDBAC-driven changes in biofilm community structure and impact on glycolipids could therefore lead to a decrease in food quality, worsened by DDBAC accumulation. Recently, Demailly et al. (2019) also demonstrated that diatom fatty acid composition can be altered by organic substances such as pesticides.

In parallel to the decrease in glycolipids noted under DDBAC contamination, we observed an increase in phospholipids. It is difficult to clearly explain this increase because the phospholipid groups analysed (i.e. PE, PC and PG) are not specific to any taxonomic group and can be found in both the plasma membranes of microalgae (Zulu et al., 2018) and in prokaryotic cells (Li-Beisson et al., 2013). One hypothesis to explain this increase could be the development of DDBAC-resistant bacteria. Indeed, the mechanism of resistance to benzalkonium chloride identified in *Pseudomonas aeruginosa* consists of an increase in the percentage of phospholipids (Sakagami et al. 1989). Considering the impact of DDBAC on biofilm bacteria would be required to determine if the increase in phospholipids observed in our study reflects the development of specific bacteria under DDBAC exposure.

Sensitivity to DDBAC differed among biofilm taxa, whose proportions changed over time, and with DDBAC concentration. This result suggests that the biofilm community acclimated to the chemical stress and that only resistant/tolerant species survived DDBAC exposure. Diatoms appeared to be minimally affected based on growth and mortality, even though the alteration of their photosynthetic structure and cell content suggests a marked impact (Figure B.2). Green algae, which experienced rapid growth in non-contaminated channels, appeared to be negatively affected by DDBAC at d10. DDBAC also had a strong effect on heterotrophic microfauna naturally present in the biofilms (e.g. rotifers), suggesting that they are highly sensitive to this contaminant. Our dataset does not allow to disentangle whether such impacts derive from direct (through water contamination) or indirect (trophic transfer from contaminated biofilms) effects. However, such results confirm the high sensitivity of aquatic microfauna and invertebrates to DDBAC, as already demonstrated for acute exposure with *Daphnia magna* (Kreuzinger et al., 2007; Leal et al., 1994; Chen et al., 2014; Lavorgna et al., 2016).

### 4.3- Effects of ALAN on the biofilm

Under our experimental conditions, ALAN did not significantly modify the structure or physiology of the autotrophic organisms in the biofilm based on the parameters that we investigated. The light intensity chosen may have been too low to generate a stress during continuous light exposure, at the timescale of the experiment.

Concerning the polar lipid classes, an increase in MGDG or DGDG is often observed during low light intensity exposures (Gushina and Harwood, 2009). In the present experiment, while light was low, the duration of exposure differed, which could explain the absence of a significant effect on these two glycolipids over time. Conversely, high light levels can lead to a decrease in polar lipids, especially phospholipids, in favour of TAGs, especially in the case of filamentous green algae such as *Cladophora* spp. (Gushina and Harwood, 2009). Changes in lipid content with ALAN were not clearly observed here, which could be due to compensation phenomena or to our averaging of the lipid mixture across different organisms present within a natural biofilm, which differs from the analysis of an isolated chlorophyte strain.

### 4.4- Combined effects of DDBAC and ALAN on the biofilm

Contrary to our hypotheses, DDBAC and ALAN exposure did not show interaction effects on any of the biological endpoints monitored in this experiment. However, ALAN impacted the fate of DDBAC by decreasing exposure concentrations in the medium. DDBAC exposure caused significant impacts on biofilm function and structure, especially impacting autotrophs. Given the wide range of uses of DDBAC, our study highlights that its transfer to the aquatic environment may result in a significant impairment of the functioning of aquatic ecosystems. As the high concentration of DDBAC tested drove most of the changes observed in biofilm composition and physiology, we cannot exclude that DDBAC exposure masked possible interactions with ALAN. Further experiments combining lower concentrations of DDBAC with ALAN would be required to completely rule out any interaction effects on aquatic biofilms.

It would be of interest to perform more in-depth lipid analyses to determine whether fatty acids show a clearer difference between ALAN treatments than it was visible from a global assessment of lipid classes. Amini Khoeyi et al. (2011), studying *Chlorella vulgaris*, showed that a prolonged photoperiod can decrease microalgal content of monounsaturated (MUFA) and polyunsaturated (PUFA) fatty acids. Such a decrease could be enhanced by DDBAC activity, which can disrupt lipid membranes and degrade lipids, thereby possibly exacerbating the negative effect of the prolonged photoperiod on MUFA and PUFA content. A decrease in essential polyunsaturated fatty acids, such as certain omega-3 and omega-6, could alter the nutritional quality of biofilms and decrease the energy supply along the trophic chain (Brett and Müller-Navarra, 1997).

## 5- Conclusion

In this experiment, we demonstrated that DDBAC exposure strongly impacts river biofilms. DDBAC accumulated quickly in biofilms and damaged the photosynthetic material, in turn altering photosynthesis. There was also evidence for an impact on the structure of the biofilm as changes in its taxonomic composition were observed. Indeed, even though the mortality index did not show greater mortality in the exposed diatoms compared with the controls, microscopic observations suggest that a marked number of individuals were strongly impacted on a physiological level (degraded cell content) and were potentially not viable. This hypothesis is supported by the results obtained during the lipidomic analysis, highlighting a strong decrease in the lipid classes associated with thylakoid membranes specific to microalgae. At elevated concentrations, we highlighted striking effects of DDBAC on the aquatic biota. As DDBAC is a common contaminant with multiple domestic uses, complementary research is needed to characterize its impacts at environmentally relevant concentrations.

In contrast, ALAN did not lead to any significant change in biofilms, even though the ALAN treatment modified their exposure to DDBAC. In view of the likely increase in DDBAC concentration in urban waters (downstream of WWTPs) where ALAN is often common, it would also be necessary to investigate whether the effects identified in the present study are manifested at environmental concentrations in order to reassess the risk posed by DDBAC to aquatic biodiversity.

## Aknowledgements

The authors aknowledge the financial support from the Institut National de Recherche pour l’Agriculture, l’alimentation et l’Environnement (INRAE) and the Groupe de Recherche Interuniversitaire en Limnologie (GRIL).

## Appendices

### A- Preliminary experiment

This preliminary experiment aimed to test the sensitivity of our biofilm to dodecyldimethylbenzylammonium (DDBAC) to select a sub-lethal concentration to be used in the main experiment. We ran a dose-response experiment in which we tested nine concentrations ranging between 17.5 µg.L^-1^ and 175 mg.L^-1^ (following a logarithmic increase) for 4 hours. Each concentration was tested twice, one test at a mean light level of 16.86 µmol.s.m^-^² and the other kept in the dark.

Glass slides (26.5 cm × 6 cm) colonized by biofilms for three months were scraped and the biofilms put into 500 mL of water. For each concentration, we contaminated 1.5 mL of the biofilm solution. After four hours of contamination, we analysed the photosynthetic efficiency of the samples with a Phyto-PAM (see section 2.3).

**Figure A.**
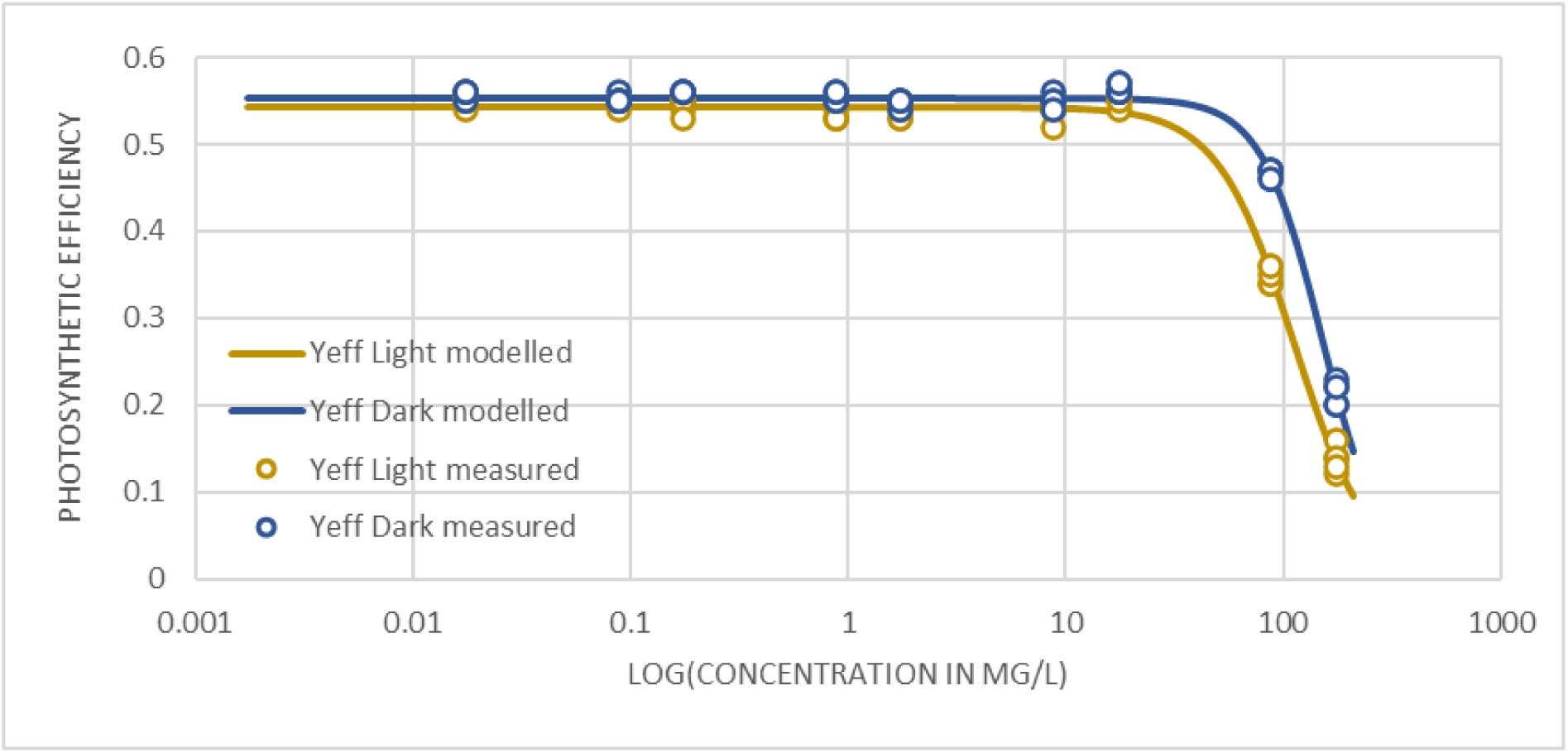
**Dose-Response curve for photosynthetic efficiency in biofilms exposed to light or dark and increasing DDBAC concentration for 4 hours. Yeff = Photosynthetic efficiency.**

The dose-response curve suggests an EC50 of 112 ± 3 mg.L^-1^ and a EC5 of 34 ± 3 mg.L^-1^ for photosynthesis inhibition in the biofilm exposed to light. For the biofilm kept in the dark, we found EC50 of 151 ± 3 mg.L^-1^ and a EC5 of 58 ± 3 mg.L^-1^. This result allowed us to choose the concentration of 30 mg.L^-1^ for the 10 days biofilm exposure which is the lowest concentration with a minimum effect on photosynthetic efficiency.

### B- Effects of DDBAC on chloroplasts

**Figure B.1.**
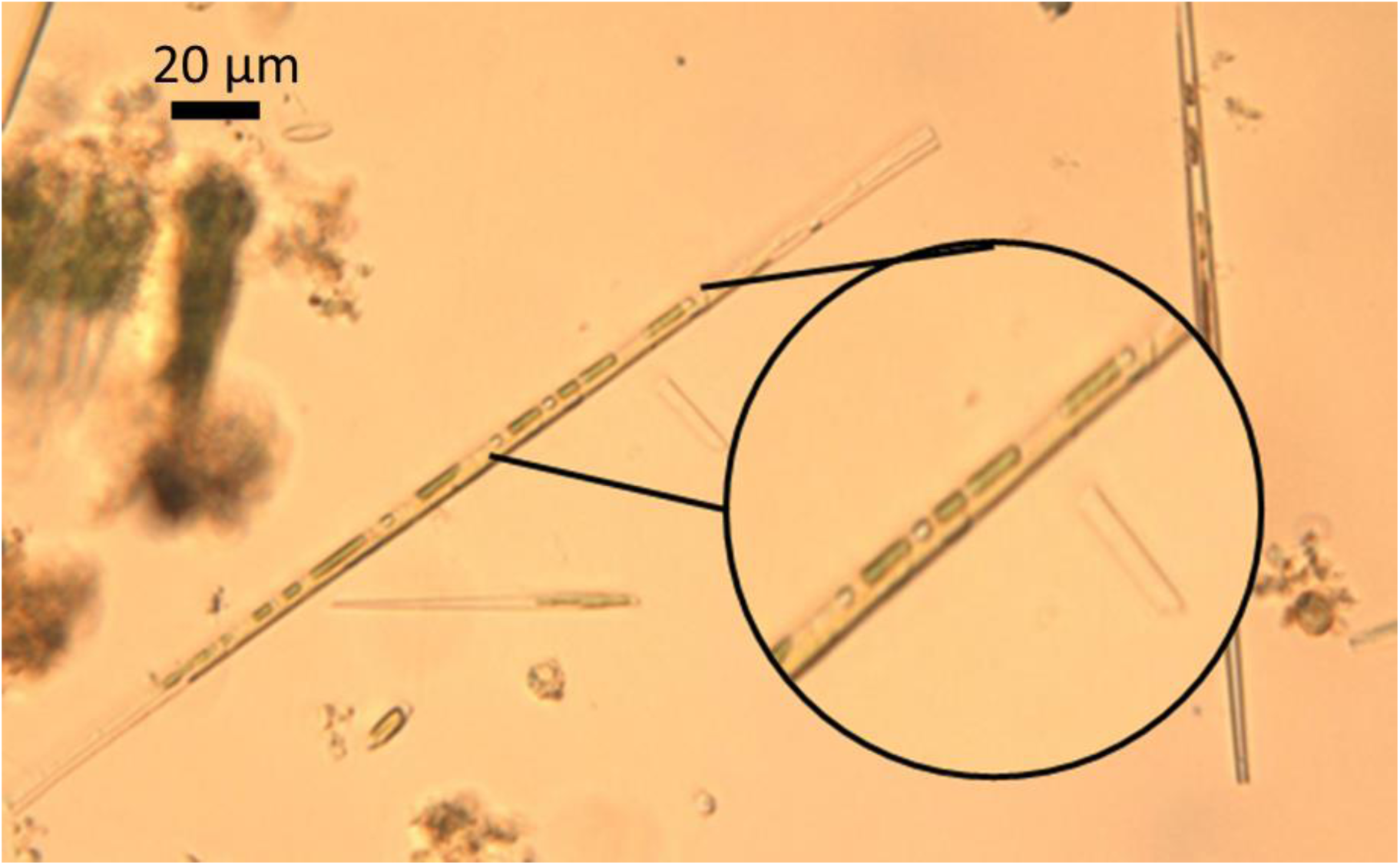
**Chloroplasts of diatoms in non-contaminated biofilm. x400 magnification**

**Figure B.2.**
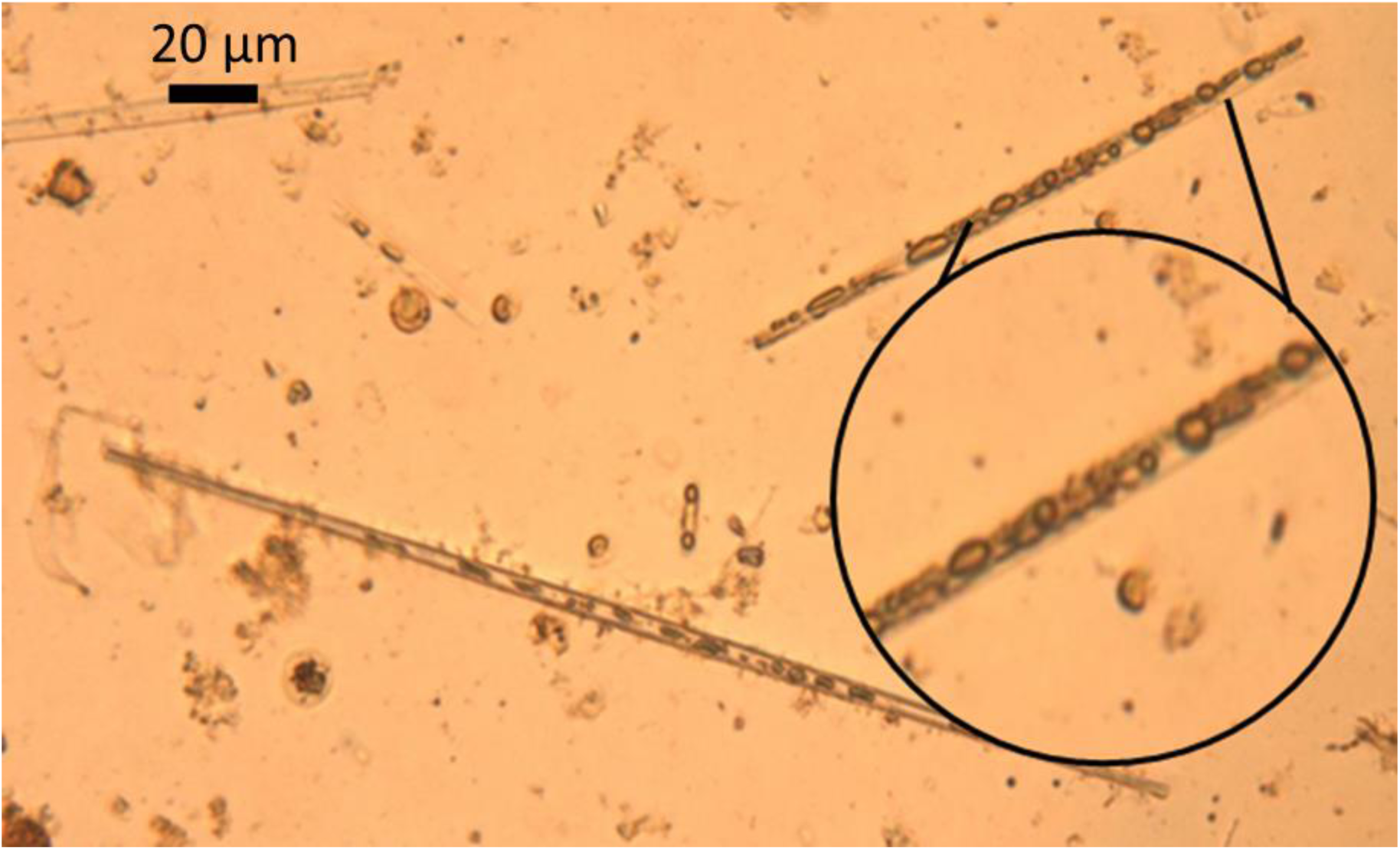
**Chloroplasts of diatoms in DDBAC contaminated biofilm after 10 days of exposure. x400 magnification.**

### C- HPLC gradients for the lipidomic analysis

**Table C.1.** HPLC gradients for phospholipid and glycolipid analysis.

| Time (min) | 40 mmol.L<sup>-1</sup> Ammonium acetate buffer (%) | Acetonitrile (%) |
| --- | --- | --- |
| 0 | 5 | 95 |
| 2 | 5 | 95 |
| 7 | 30 | 70 |
| 10 | 30 | 70 |
| 11 | 5 | 95 |
| 13.7 | 5 | 95 |

**Table C.2.** HPLC gradients for triglyceride analysis.

| Time (min) | Solvent A* (%) | Solvent B* (%) |
| --- | --- | --- |
| 0 | 50 | 50 |
| 0.3 | 50 | 50 |
| 5.3 | 1 | 99 |
| 7.3 | 1 | 99 |
| 8.3 | 50 | 50 |
| 9.8 | 50 | 50 |

\* Solvent A: solution of acetonitrile/water/40 mmol.L<sup>-1</sup> ammonium acetate buffer (600/390/10, v/v/v)
Solvent B: solution of isopropanol/acetonitrile/1 mol.L<sup>-1</sup> ammonium acetate buffer (900/90/10, v/v/v)

### D- Lipid standards

Polar lipid standards were purchased from Avanti Polar Lipids. Quantitations of phosphatidylcholine (PC), phosphatidylethanolamine (PE) and phosphatidylglycerol (PG) were respectively carried out with 1-palmitoyl-2-oleoyl-glycero-3-phosphocholine or PC (16:0/18:1) (850457), 1-palmitoyl-2-oleoyl-sn-glycero-3-phosphoethanolamine or PE (16:0/18:1) (850757), and 1-palmitoyl-2-oleoyl-sn-glycero-3-phospho-(1’-rac-glycerol) or PG (16:0/18:1) (840457).

For glycolipids, monogalactosyldiacylglycerol (840523), digalactosyldiacylglycerol (840524) and sulfoquinovosyldiacylglycerol (840525) from plant extracts were used as standards. Quantitation was performed with the following molecular species: MGDG (16:3_18:3) (63% of the total MGDG standard), DGDG (18:3_18:3) (22% of the total MGDG standard), and SQDG (34:3) (78% of the total MGDG standard).

1,2-diheptadecanoyl-sn-glycero-3-phosphocholine or PC (2×17:0) (850360) was used as internal standard for PC phospholipids, 1,2-diheptadecanoyl-sn-glycero-3-phosphoethanolamine or PE (2×17:0) (830756) was used as internal standard for PE phospholipids, and 1,2-diheptadecanoyl-sn-glycero-3-phospho-(1’-rac-glycerol) or PG (2×17:0) (830456) was used as internal standard for PG phospholipids, and both MGDG, DGDG and SQDG glycolipids.

1,2-dipalmitoyl-sn-glycero-3-O-4’-(N,N,N-trimethyl)-homoserine or DGTS (2×16:0) (857464) was used for the diacylglyceryltrimethylhomo-Ser (DGTS) lipids. 1,2-dipalmitoyl-sn-glycero-3-O-4’-[N,N,N-trimethyl(d9)]-homoserine or DGTS-d9 (2×16:0) (857463) was used as internal standard for DGTS lipids

Triglycerides were purchased from Sigma-Aldrich. Tristearin or TAG (3×18:0) (≥99%, T5016) was used as the calibration standard while TAG (3×17:0) (≥99%, T2151) was used as the internal standard.

### E- Mass spectrometry parameters for lipid analysis

**Table E.1.** Mass spectrometry parameters for phospholipid and glycolipid analysis.

|  | Curtain gas | CAD | IonSpray | Temperature | Ion source gas 1 | Ion source gas 2 | Declustering potential | Collision energy |
| --- | --- | --- | --- | --- | --- | --- | --- | --- |
| MGDG | 30 psi | 3 | -4500 V | 450°C | 30 psi | 60 psi | -61 V | -28 V |
| DGDG | 30 psi | 3 | -4500 V | 450°C | 30 psi | 60 psi | -61 V | -28 V |
| SQDG | 30 psi | 3 | -4500 V | 450°C | 30 psi | 60 psi | -126 V | -66 V |
| PE | 30 psi | 3 | -4500 V | 450°C | 30 psi | 60 psi | -50 V | -50 V |
| PG | 30 psi | 3 | -4500 V | 450°C | 30 psi | 60 psi | -100 V | -50 V |
| PC | 30 psi | 3 | -4500 V | 450°C | 30 psi | 60 psi | -100 V | -50 V |

**Table E.2.**
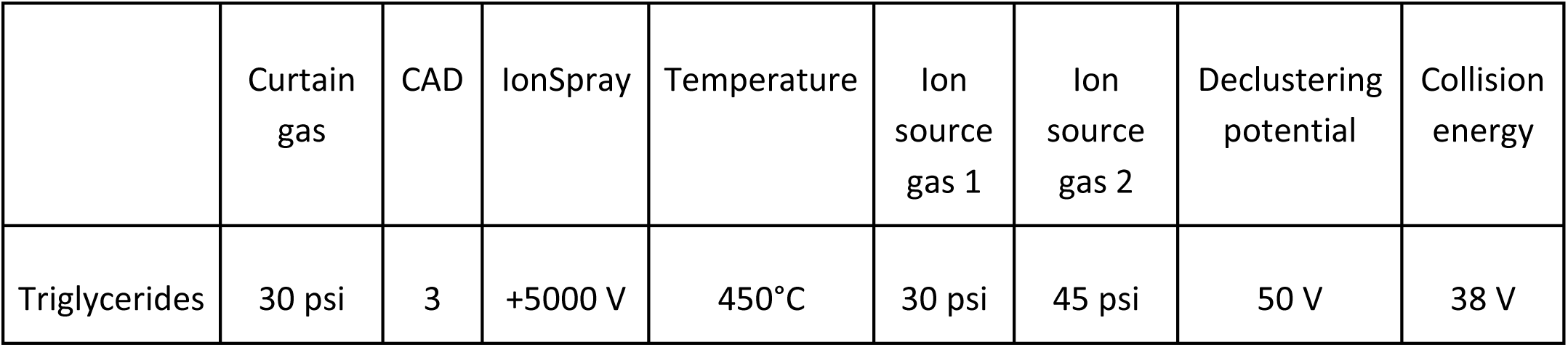
Mass spectrometry parameters for triglyceride analysis.

|  | Curtain gas | CAD | IonSpray | Temperature | Ion source gas 1 | Ion source gas 2 | Declustering potential | Collision energy |
| --- | --- | --- | --- | --- | --- | --- | --- | --- |
| Triglycerides | 30 psi | 3 | +5000 V | 450°C | 30 psi | 45 psi | 50 V | 38 V |

